# Incorporation of a cost of deliberation time in perceptual decision making

**DOI:** 10.1101/2024.01.31.578067

**Authors:** Shinichiro Kira, Ariel Zylberberg, Michael N. Shadlen

## Abstract

Many decisions benefit from the accumulation of evidence obtained sequentially over time. In such circumstances, the decision maker must balance speed against accuracy, and the nature of this tradeoff mediates competing desiderata and costs, especially those associated with the passage of time. A neural mechanism to achieve this balance is to accumulate evidence in suitable units and to terminate the deliberation when enough evidence has accrued. To accommodate time costs, it has been hypothesized that the criterion to terminate a decision may become lax as a function of time. Here we tested this hypothesis by manipulating the cost of time in a perceptual choice-reaction time task. Participants discriminated the direction of motion in a dynamic random-dot display, which varied in difficulty across trials. After each trial, they received feedback in the form of points based on whether they made a correct or erroneous choice. They were instructed to maximize their points per unit of time. Unbeknownst to the participants, halfway through the experiment, we increased the time pressure by canceling a small fraction of trials if they had not made a decision by a provisional deadline. Although the manipulation canceled less than 5% of trials, it induced the participants to make faster decisions while lowering their decision accuracy. The pattern of choices and reaction times were explained by bounded drift-diffusion. In all phases of the experiment, stopping bounds were found to decline as a function of time, consistent with the optimal solution, and this decline was exaggerated in response to the time-cost manipulation.

## Introduction

A decision is a commitment to a proposition or plan based on evidence from the environment or memory. Often a decision maker is allowed the opportunity to deliberate over a sequence of samples of evidence, which bear on the proposition at hand. In that case, the decision is not just about the proposition but also about when to stop deliberating. Such sequential sampling with optional stopping invites a policy that would balance the speed of making a decision against accuracy. For decisions about a perceptual category (e.g., present/absent or left/right), sequential sampling can be viewed as a natural extension of signal detection theory (Green and Swets, 1966; Luce, 1986; Gold and Shadlen, 2007; Shadlen and Kiani, 2013). In this theory, a criterion on evidence is viewed as a policy that instantiates a negotiation between the value (or cost) of answering one way or another, correctly or incorrectly. The same can be said of sequential sampling, with the addition that time, or the number of samples, is also part of the negotiation.

For many decisions, it is reasonable to presume that samples of evidence are conditionally independent and identically distributed. That is, given that the stimulus is in a particular state (e.g., present or absent; left or right), the samples of evidence are like random draws from some distribution of numbers. The samples are informative to the extent that the distributions associated with each hypothesis, state, or alternative are distinct. In that case, a sensible strategy is to accumulate evidence in units of the logarithm of the likelihood ratio (logLR) that the evidence would have been observed under one hypothesis or its alternative(s) (Wald, 1947; Wald and Wolfowitz, 1948; Laming, 1968; Link, 1992). The deliberation terminates when the logLR equals (or exceeds) a criterion level or bound. An important insight from signal detection theory is that the evidence need not be in units of logLR, *per se*. Any monotonic transformation, such as conversion to spike rate, is sufficient to allow a single criterion to achieve the “negotiation” mentioned above (Green and Swets, 1966; Kira et al., 2015). However, the matter is more complex when the reliability of the evidence bearing on the decision is unknown.

If decisions are to be made about stimuli that differ in difficulty, as is often the case in tests of perception, the bound may need to be adjusted for each type of stimulus. More importantly, when costs are incurred as a function of decision time (e.g., to maximize reward rate), it may be sensible for the terminating bound to be time-dependent (Gershman et al., 2015). In particular, it has been shown that if the level of difficulty is not known to the decision-maker at the beginning of deliberation, then in order to maximize the reward rate, the magnitude of the stopping bound should decline as a function of time (Frazier and Yu, 2008; Drugowitsch et al., 2012). We refer to this decline as a collapse of the decision bounds. There is experimental evidence for an equivalent mechanism shown in neurophysiological studies of perceptual decision-making in monkeys (Hanks et al., 2014). It is unknown, however, what environmental cues or experiences with a task, lead a decision-maker to implement such adjustments to their stopping policy.

We wished to ascertain whether human participants would adjust their termination criteria to accommodate a subtle manipulation of the cost of deliberation time. We used a variant of a direction discrimination task for which it is well established that choices and reaction times (RT) conform to the sequential sampling models. We established conditions that encouraged participants to optimize the number of correct versus error choices per time and then surreptitiously manipulated time costs. We developed a novel computational method to estimate the time-course of decision termination bounds from the pattern of choice and RTs. The method reveals that each participant approximated the optimal performance with their own distinct shape of a collapsing bound. All participants changed their behavior in response to the subtle experimental manipulation by adjusting the starting height of the bound and its rate of collapse, while preserving their distinct shape.

## Results

### A manipulation affecting the time cost of deliberation

Four human participants performed a reaction-time version of the random dot motion discrimination task (Fig. 1a). On each trial, the motion coherence was chosen at random from the set {±0, ±3.2, ±6.4, ±12.8, ±25.6, ±51.2%}, where the sign denotes the direction (negative for leftward and positive for rightward). We refer to the absolute value of coherence as the *motion strength* (see Methods). When the participants reached decisions, they indicated their choice by making an eye movement to the right or left peripheral target. Participants were incentivized to make appropriate speed-accuracy tradeoff based on scoring of their performance. They earned one point for a correct choice and lost one point for an erroneous choice. The score did not change if they failed to maintain fixation or made an inaccurate eye movement to the target. When these events occurred, the ongoing trial stopped immediately and a next trial followed. After each trial, in addition to auditory feedback on the accuracy of the choice, they received visual feedback and a tally of the current point total as well as an intuitive graphic that displayed their points per minute (*earning rate*, from here on) (Fig. 1a). Note that random guesses do not improve the earning rate because the expected score for a random choice is 0. Instead, they could increase the rate by answering quickly and accurately with saccades and by maintaining accurate gaze fixation, since inaccurate fixation would add time to the experiment, thereby lowering the earning rate.

**Figure 1.**
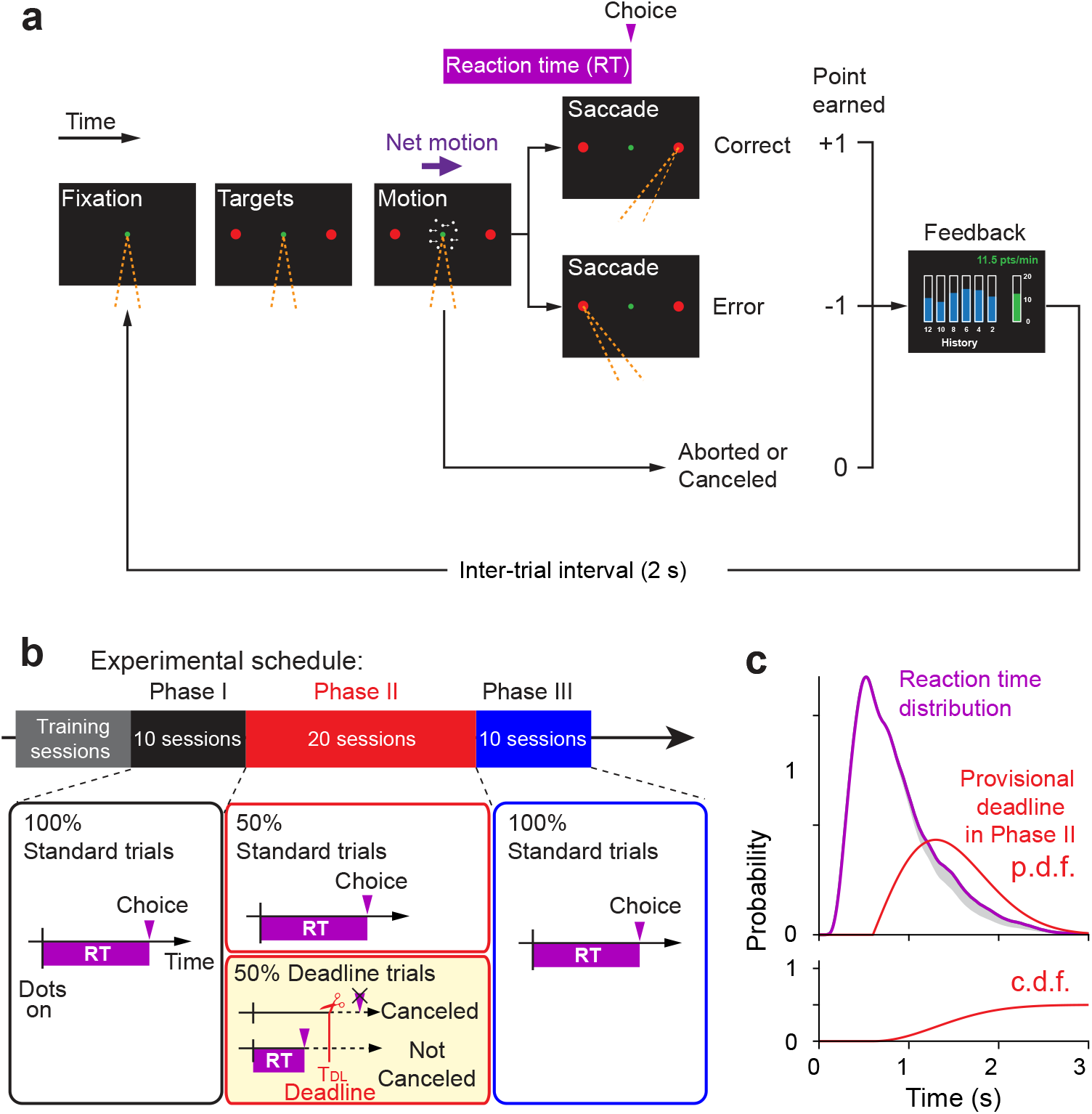
Experimental design. Participants made perceptual decisions in a choice-reaction time design. Unbeknownst to the participants, the sessions were organized in three phases. (a) Random-dot motion (RDM) task. Participants initiate a trial by fixating a central spot (green). Two peripheral targets then appear (red circles). After a variable delay, the RDM appears in a virtual aperture centered on the fixation point. Participants must decide the net direction of motion (right or left) and report this decision by making an eye movement to the choice-target (red) in the corresponding direction. Participants earn 1 point for a correct decision and lose 1 point for an incorrect decision. The participant receives immediate auditory feedback for correct and erroneous choices as well as visual feedback about their points earning rate over the preceding two minutes (green bar) and its history during the session (blue bars). Trials aborted by blinks, poor fixation, or stochastic cancellation (Phase II only) incur no change in points but affect the points earning rate. Orange dashed lines indicate the gaze direction. (b) Experimental schedule. The participants performed training sessions until their point-earning rate was stable, followed by the three phases of the experiment. Phase I comprised 10 sessions identical to the training. Phase II comprised 20 sessions in which half of trials (randomly interleaved) were assigned provisional deadlines (yellow box). Phase III comprised 10 sessions identical to Phase I (i.e., without provisional deadlines). (c) *top*, Reaction time distribution from Phase I (purple) and the distribution of provisional deadlines planned for Phase II (red) for one participant (participant A). The reaction time distribution was smoothed by Epanechnikov kernel (inverted parabola) with s.d. = 0.1 s. *bottom*, cumulative probability density function (c.d.f.) of provisional deadlines. The deadlines are planned surreptitiously on half the trials, hence the area under the p.d.f. (and upper asymptote of the c.d.f.) is 0.5. Shaded gray area shows 10% of RT distribution expected to be canceled by deadlines without a change in decision strategy from Phase I to II.

The study was conducted in three phases (Fig. 1b). In Phase I, participants received extensive training until they achieved a stable earning rate. During training sessions, participants were acclimated to the game-like structure of the task. Each session lasted 13 minutes. Participants completed two sessions per day with a short break in between. They were specifically instructed to maximize their earning rate. We used the last 10 sessions to establish a baseline level of performance (2154–2389 trials; 224.4 ± 10.9 trials/session, mean ± s.d., n = 40 sessions from 4 participants). In Phase II, unbeknownst to the participant, we imposed an additional time cost. On a random half of the trials, we chose a *provisional deadline, T*_*DL*_. If the participant deliberated beyond *T*_*DL*_, the trial was canceled with termination (Fig. 1b, yellow box). From the participant’s perspective, this occurrence was not dissimilar to other aborted trials, including those with an eye blink or an inaccurate fixation (Methods). *T*_*DL*_ was drawn from a Rayleigh distribution, bespoke to each participant. The parameters of the distribution were chosen to ensure that these provisional deadlines would come to fruition on approximately 10% of trials, absent a change in decision strategy (Fig. 1c). The other half of trials had no provisional deadline, and thus did not differ from trials in Phase I. In Phase III we returned to the original paradigm without the deadlines. The phases were neither identified nor signaled to the participants. From their perspective they were playing the same game for points.

The manipulation in Phase II affected task performance. All four participants reduced their reaction times substantially, especially on the trials with low motion strength (Fig. 2a). For example, RTs decreased by 34.0 ± 7.5% at zero motion strength (mean ± s.e.m. across 4 participants), while the decrease in RTs was only 8.6 ± 6.3% at the highest motion strength (Fig. 2a). The reduction in RT from Phase I to Phase II held for both correct and erroneous choices (dark and light symbols, *p* < 10^−62^s and *p* < 10^−19^, respectively, linear regression; Eq. 2). As we shall see, this suggests a change in the time dependent criterion for terminating decisions. The increase in speed led the participants to beat the clock, so to speak. Instead of the 10% of trials that should have been terminated at *T*_*DL*_, less than 5% of trials were terminated by the provisional deadline (Table 1).

**Table 1.** Sampling distribution for provisional deadlines and the fraction of trials canceled or aborted.

| participant | Sampling distribution for provisional deadline |  |  |  | Canceled trials (%)<br>in Phase II<br>(n = 20 sessions) | Aborted trials (%) |  |  |
| --- | --- | --- | --- | --- | --- | --- | --- | --- |
| | to (s) | $\sigma_{DL}$ | mean (s) | s.d. (s) | | Phase I<br>(n = 10 sessions) | Phase II<br>(n = 20 sessions) | Phase III<br>(n = 10 sessions) |
| A | 0.6 | 0.7 | 1.48 | 0.46 | 1.4 ± 0.9 | 1.7 ± 1.4 | 2.2 ± 1.0 | 3.2 ± 1.2 |
| B | 0.3 | 0.6 | 1.05 | 0.39 | 4.3 ± 1.9 | 3.4 ± 1.2 | 2.5 ± 0.9 | 3.7 ± 2.0 |
| C | 0.6 | 1.0 | 1.85 | 0.66 | 3.6 ± 1.1 | 2.5 ± 1.3 | 2.7 ± 2.5 | 5.2 ± 2.9 |
| D | 0.6 | 0.6 | 1.35 | 0.39 | 3.7 ± 2.1 | 0.6 ± 0.5 | 1.3 ± 1.1 | 1.1 ± 0.9 |

**Figure 2.**
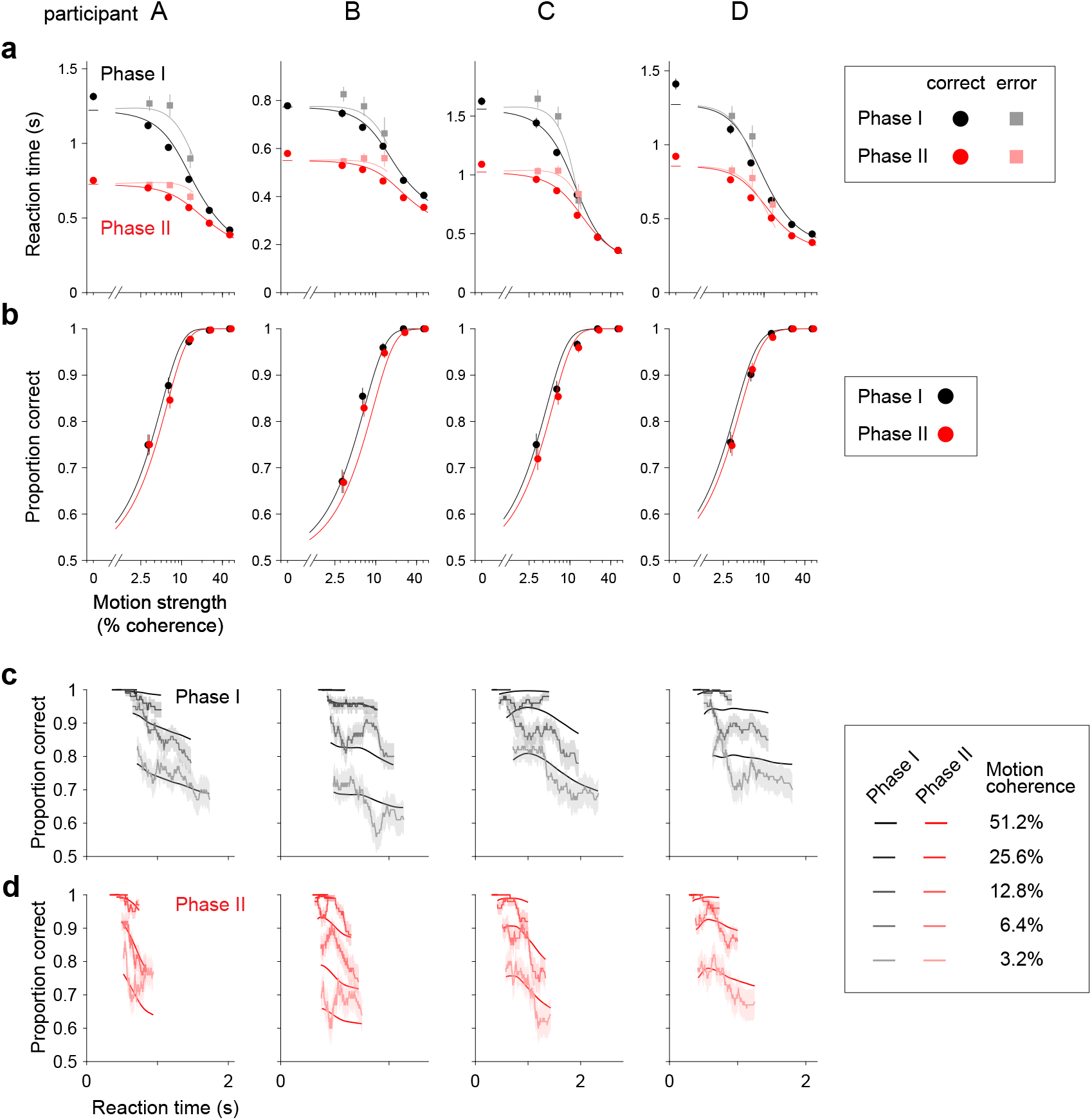
Effects of increased time cost on reaction time and accuracy for individual participants. (a) Mean reaction time for correct (circles) and error (squares) trials as a function of motion strength. Columns correspond to the four participants. Data from phases I and II are shown in black and red, respectively. Mean RTs for error trials are not shown if the participant had fewer than five error trials for a given motion strength. Error bars show s.e.m. The curves are fits of the nonparametric-bound Drift-Diffusion Model (*npb-DDM*; see Figure 3 and Methods). (b) Proportions of correct choices as a function of motion strength for Phase I (black) and II (red). Error bars show s.e.m. Curves represent the proportion of correct trials predicted by *npb-DDM*. (c) Proportion of correct choices as a function of RT, for trials from Phase I. The proportion correct was calculated independently for each motion strength, in sliding windows of 100 trials. Shading indicates s.e.m. The curves are fits of the *npb-DDM*. (d) As panel (c), for trials from Phase II. (e) Only trials without a provisional deadline are included in the Phase II data in panels (a, b, d).

The manipulation also affected decision accuracy. For each participant, accuracy was lower in Phase II than in Phase I, even when restricting the comparison to trials without provisional deadlines (Fig. 2b). We confirmed this qualitative observation by fitting a logistic regression model relating motion strength to choice accuracy (Eq. 3). The model, fit to Phase I and II trials together, includes an indicator variable to identify Phase II trials. For all participants, the regression coefficient associated with this indicator variable (*β*_Phase_; Eq. 3) is negative, indicating that accuracy was lower in Phase II. Although statistically reliable, the change in accuracy is subtle (Fig. 2b), achieving significance only in grouped data from the four participants (p<0.034; ℋ_0_ : *β*_Phase_ = 0; Eq. 4). Overall, the changes in behavior between Phases I and II suggest a change in the tradeoff between decision speed and accuracy.

An explanation of the behavioral effects of the manipulation stems from the analysis of the time-dependent accuracy functions. For Phase I trials with the same motion strength, faster decisions were, on average, more accurate (Fig. 2c). Such a decay in the time-dependent accuracy function may appear perplexing because slower decisions are thought to involve more deliberation, hence greater accuracy. What is missing from this intuition, however, is that the decision-maker is controlling the duration of the decision by exercising a stopping policy. The decrease in accuracy with slower decisions is a sign that the stopping criterion is more lax (i.e., that decision bounds *collapse* over time). The time-dependent accuracy functions in Phase II showed a more pronounced decay than in Phase I (Fig. 2d), suggesting that the behavioral adjustments in Phase II may be mediated by lower and/or faster-collapsing bounds—an idea that we formalize below with a novel variant of the drift-diffusion model.

### The speed-accuracy regime is controlled by the adjustment of the decision termination bounds

To investigate which aspects of the decision policy changed from Phases I to II, we fit the data to a bounded evidence-accumulation model. In the model, the decision follows a driftdiffusion process with a constant diffusion coefficient and a drift-rate proportional to the motion coherence (Fig. 3a). The decision process ends when the accumulated evidence (termed the *decision variable*) crosses an upper or lower bound at ±*B*(*t*) signaling a rightward or leftward choice, respectively. The reaction time comprises the time it takes for the diffusing particle to reach one of the bounds (the decision time, *T*_*d*_) plus the non-decision time (*T*_*nd*_) that groups sensory and motor latencies, which are assumed independent of decision difficulty. Given the parameters of a DDM, the probability of decision termination at any moment can be computed by propagating the unabsorbed states of the decision variable over time and recording the probability mass that is absorbed at the upper or lower bound (Fig. 3b). Our working hypothesis is that the changes in the speed-accuracy tradeoff from Phases I to II are mainly explained by adjustments in the decision termination bound, *B*(*t*).

**Figure 3.**
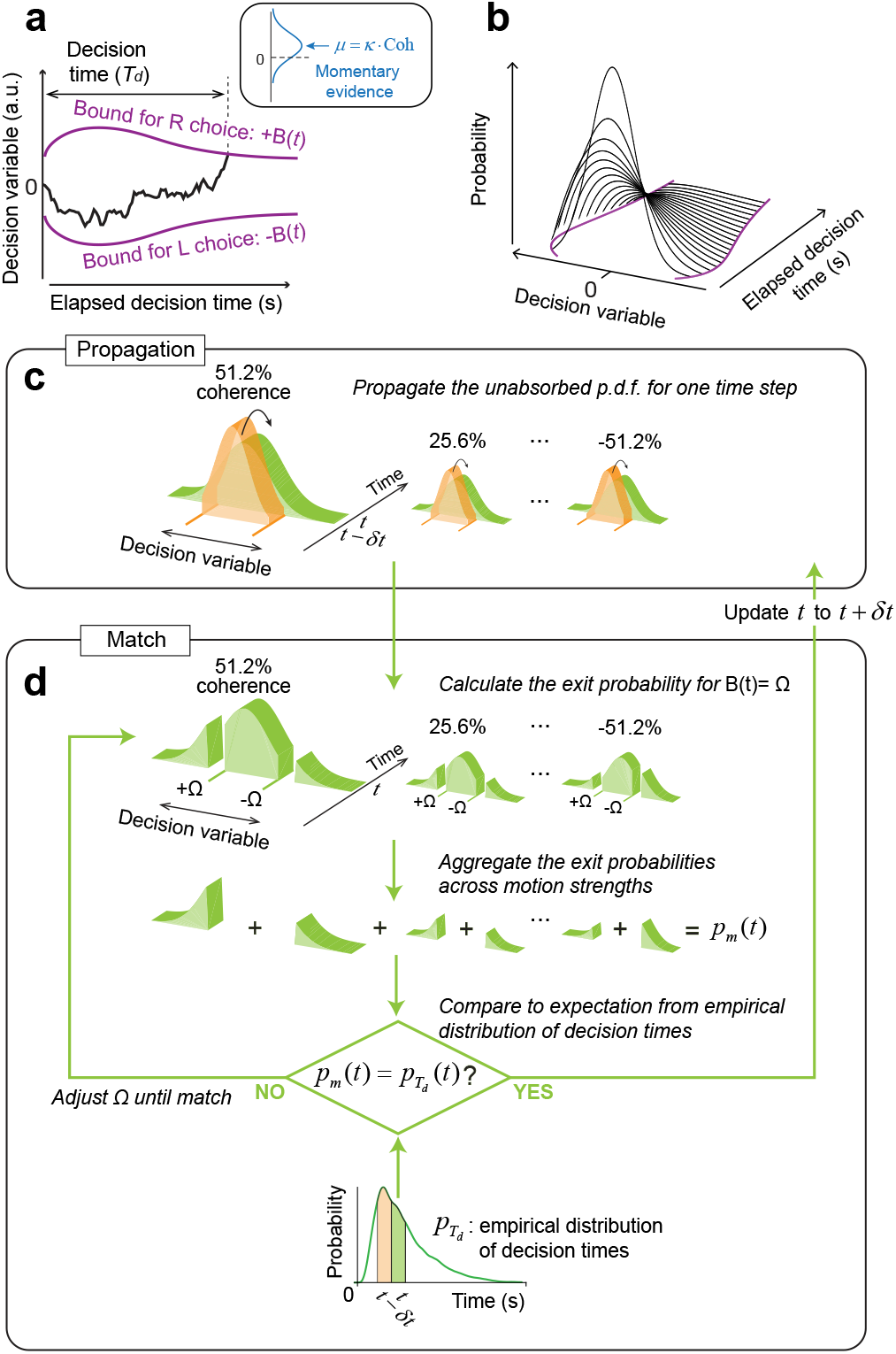
Bounded evidence-accumulation model. (a) Schematic representation of the drift-diffusion model with time-dependent bounds. The decision variable, *x*(*t*), represents the accumulation of noisy momentary motion evidence over time. It follows a drift-diffusion process where the drift rate is proportional to the motion coherence (inset). The bounds are symmetric about the origin and vary as a function of time (purple curves). Reaching a bound determines the choice and decision time (*T*_*d*_) simultaneously. The observed reaction time is the sum of *T*_*d*_and and the non-decision latencies, *T*_*nd*_(not shown). Only *T*_*d*_depends on the direction and strength of motion. (b) If the decision bounds were known, the probability of terminating the decision at each moment in time (referred to as the *exit* probability) can be calculated iteratively. This involves propagating over time the probability of the state of *x*(*t*) (black curves) that remains unabsorbed by the bounds (purple curves) and measuring the probability mass that has been absorbed. (c-d) Schematic illustration of the algorithm used to estimate the decision bounds, *B*(*t*). The algorithm proceeds iteratively over time, alternating between *propagation* of the state of the decision variable, *x*(*t*), and *match* to the exit probabilities inferred from the observed reaction times. (c) The probability density function (*pdf*) for the state of the decision variable is propagated for one time-step: from time *t*−*δt* (orange) to *t* (green). This step is performed independently for each motion coherence. (d) The *match* step establishes the height of the bound at time *t* that matches the probability of decision termination at time *t* between the model and the data. We calculate the exit probability for each motion coherence assuming that the bound at time *t* is equal to Ω (top row of panel). Exit probabilities are aggregated across motion coherences (second row). The total exit probability, *p*_*m*_(*t*) (*m* for model), is compared to the value expected from the empirical distribution of reaction times, 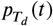. We compute 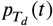 from an estimate of decision times obtained by subtracting the expected non-decision time, *μ*_*nd*_, from each trial’s reaction time. We fit the resulting decision time estimates with a nonparametric kernel density function, from which we calculate 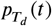 (bottom row). We find the value of Ω by root-finding until *p*_*m*_(*t*) equals 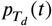. Then time is increased by *δt* and the process is repeated until all the probability mass has been absorbed.

To test this hypothesis, we developed a method that allows us to derive the shape of the decision-termination bounds without assuming their functional form. We call this model the nonparametric-bound Drift-Diffusion Model (*npb-DDM*) (see also Glickman et al., 2022). The key intuition behind this model is that for a given signal-to-noise, the distribution of decision times is uniquely determined by the shape of the decision bounds; we reverse the logic and use the distribution of empirical decision times to infer the shape of the time-varying decision bounds without fitting. Assuming that the variability of the non-decision times is small compared to that of the decision times (e.g., Zylberberg et al., 2016), the distribution of decision times, 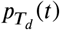, can be approximated by the distribution of reaction times shifted in time by the mean non-decision time. The derivation of the bounds involves two steps, *propagation* and *match*. We assume that the state of the decision variable at time *t* = 0 is a delta function centered at *x* = 0 (no accumulated evidence for either choice). In the *propagation* step, the probability distribution for the state of the decision variable is propagated forward in time for a single time step, from *t* − *δt* to *t*. The propagation is performed independently for each motion coherence (Fig. 3c). In the *match* step, we determine where the bound should be placed at time *t*, such that the probability of crossing the upper or lower bound, when aggregated across motion coherences, is equal to the probability of making a decision at time *t* obtained from the empirical distribution of decision times, 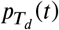 (Fig. 3d). We iterate the *propagation* and *match* steps over time until all the probability mass has been absorbed at one of the decision bounds. The *npb-DDM* allows us to fit single-trial choice and reaction time data with only three parameters: a signal-to-noise parameter (*κ*) and two parameters for the distribution of non-decision times (see Methods). The *npb-DDM* provides an excellent fit to reaction times and accuracy for each phase and participant (solid lines in Fig. 2, Fig. S1), and recovers the true shape of bounds when these are known in simulations (Fig. S2).

### Bound adjustments are consistent with earning rate maximization

The fitted models show that the participants decreased their decision-stopping criterion (|B(t)|), to accommodate the increased time cost in Phase II. Fig. 4a shows the shape of the bounds inferred by the model for Phases I (black) and II (red). For all participants, the bounds were closer to zero in Phase II than in Phase I. The change in bounds accounts for almost all of the RT reduction from Phase I to II, while changes in other model components (drift rate or non-decision time) account for only a small reduction (Fig. S3). The lowered bound height primarily reduced the proportion of trials with long RTs, explaining a more pronounced reduction in the RT for lower coherence trials (Fig. 2a). In contrast, a decrease in the decision accuracy was moderate for lower coherence trials because the accuracy in low coherence trials was close to, or at, chance level even during Phase I (Fig. 2b). Therefore, the pattern of bound adjustment from phases I to II is sensible as it speeds up what would have been low-accuracy long-RT trials.

**Figure 4.**
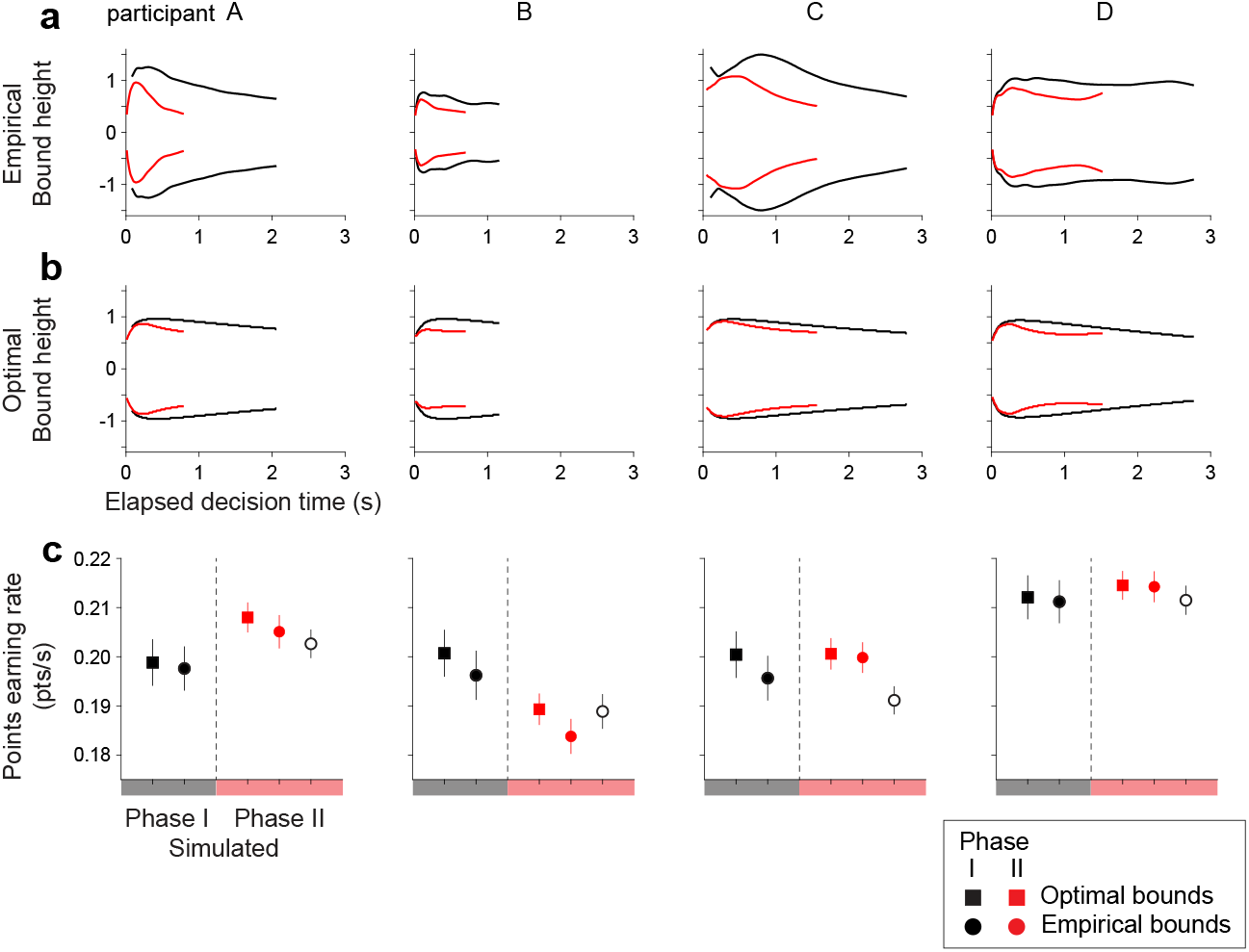
Adjusted decision termination bounds under increased time costs. (a) Shape of the decision-termination bounds derived with the *npb-DDM*. Bound shapes for Phases I and II are shown in black and red, respectively. The bound shapes were derived from observed RTs (see Fig. 3). The bounds are shown for the range corresponding to the 1^st^–99^th^ percentile of decision times for each participant and phase. Columns correspond to the four participants. (b) Shape of the decision-termination bounds that would maximize the reward rate. Optimal bounds for Phase I and II are shown in black and red, respectively. The bound shapes were derived using Dynamic Programming to solve the Bellman equations. Signal-to-noise (*κ*) and mean non-decision time (*μ*_*nd*_) were fixed at the values obtained from the *npb-DDM* fits for each participant. (c) Simulated points earning rate in Phase I and II with optimal and fitted bounds. Symbols and error bars show mean ± s.e.m. of points earning rate in each Phase (10 sessions in Phase I or 20 sessions in Phase II). Open symbols show the expected points earning rate if the decision-termination bounds from Phase I were used in Phase II without adjustment.

We used dynamic programming to determine whether the adjustment of the bounds approximates the adjustment expected under the optimal policy. In this framework, a decision policy is a deterministic mapping from a state of accumulated evidence and time, to an action. It can be succinctly represented by bounds that separate the regions where it is favorable to continue gathering evidence from those where it is favorable to make a rightward/leftward choice. The optimal policy was derived independently for each participant and each phase of the task, using the parameters obtained from the fits of the *npb-DDM*. The optimal strategy is to set the bounds lower in Phase II (Fig. 4b), akin to what we obtained from the fits of the *npb-DDM* (Fig. 4a). Based on our simulation (Fig. 4c), three out of four participants (A, C, D) would have experienced a lower earning rate had they carried their bounds from Phase I to II without adjustment (filled red circle vs. open black circle in Fig. 4c). Overall, the participants achieved 99% of the optimal points earning rate in each Phase (99.2 ± 0.7% in Phase I, 99.2 ± 0.7% in Phase II, n = 4 participants; Fig. 4c), supporting the notion that they sensibly adapted the decision bounds to the time cost manipulation.

### Bound shape is idiosyncratic

The shape of the decision bounds derived from the *npb-DDM* fits do not seem to have a simple parameterization (e.g., constant or exponential) (Fig. 4a). Therefore, it is unclear how many degrees of freedom (parameters) the brain may need to control in order to establish the decision bound. More parameters would provide more flexibility, but at a cost—that is, it would take many trials to find the right combination. We hypothesize that bound shapes are idiosyncratic (i.e., characteristic of each participant), and that the bounds are adjusted to different speed-accuracy settings by simple shape-preserving transformations. To test this hypothesis, we used the bound shape obtained from Phase I of the task to fit the data of Phase II. We allowed the bounds obtained from Phase I to change by simple multiplicative scaling in time and/or magnitude (Fig. 5a). This procedure was performed within each participant or between participants; Phase I bounds of one participant were scaled to fit Phase II data of the same participant or a different participant (Fig. 5b). The scaled bounds tended to explain the behavior of the same participant better than those of the others (p=0.02, see Methods). While this result should be considered preliminary given the small number of participants, it suggests that the termination criterion of a decision (and hence, also the distribution of reaction times and confidence judgments) may be an individual trait (Ais et al., 2016).

**Figure 5.**
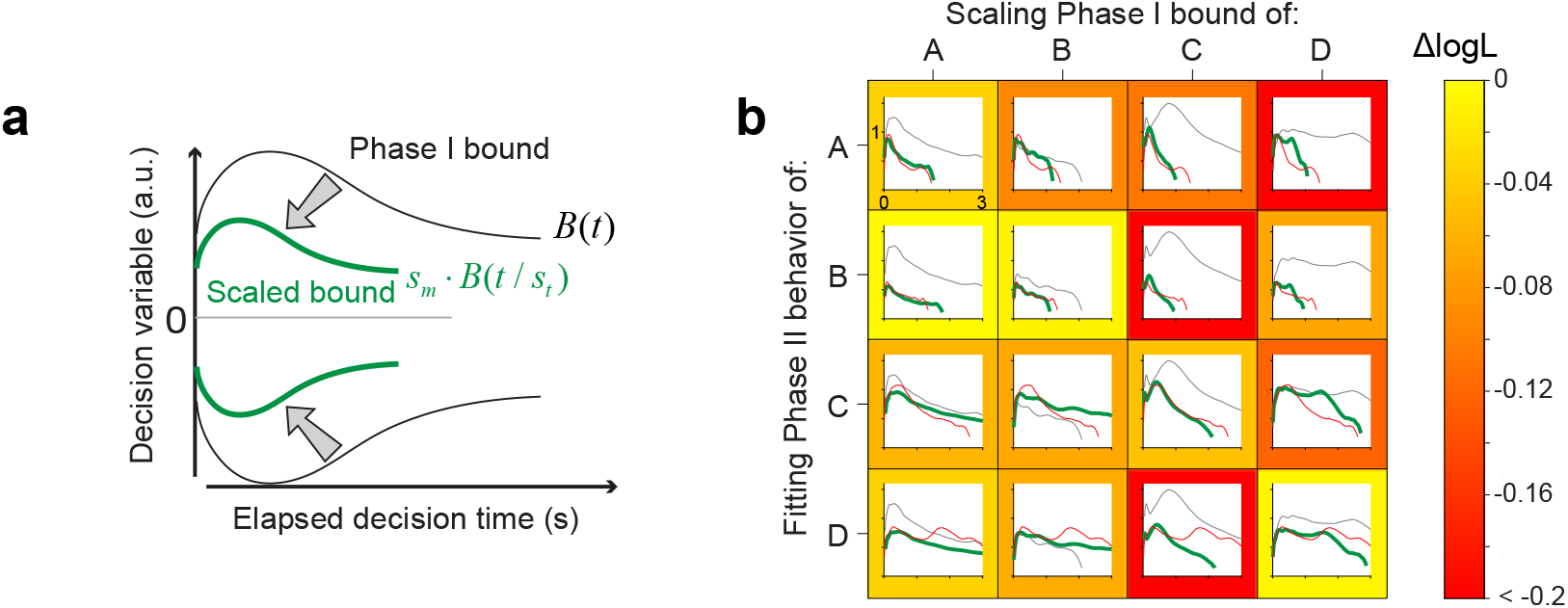
Scaling of Phase I decision bound to fit Phase II data. (a) Schematic illustration of the transformation method. The bounds obtained from Phase I (black curves) are scaled linearly as a function of magnitude and time to fit the data from Phase II (only trials without a provisional deadline). The scaling parameters (*s*_*t*_ and *s*_*m*_) were fit to the choice and RT data from Phase II. (b) Distinct bound profiles associated with individual participants. The grid shows the degree to which the Phase-I decision bound (black curve) for participant *i* (column) can be transformed to explain the Phase II data for participant *j* (row). The thick green curve indicates the best-fitting decision bounds obtained with the bound scaling approach. The red curve shows the decision bounds derived from fitting the *npb-DDM* to the Phase II data for participant *j* (row). Note that the green and red curves tend to be more similar when the same participant’s data are used for scaling and fitting (grids along the diagonal). The frame color of each grid reflects the quality of the fit, quantified as the difference in log-likelihood per trial (ΔlogL; color bar) between the model that uses the bound-scaling approach and the *npb-DDM* fit to Phase II.

### Rapid adjustment of decision policy after experiencing a cancellation

Our task requires adjusting the decision policy over time based on scattered unpredicted signals—here the trial cancellations. To analyze how canceled trials affected subsequent decisions, we compared the reaction times before and after canceled trials. Reaction times were reduced immediately after a trial cancellation (p<0.05, two-sample t-test comparing RTs on the last two trials before and the first two trials after a canceled trial) (Fig. 6a). These adjustments occurred even with limited experience of trial cancellations, as three out of four participants markedly increased the decision speed even within the first session of Phase II (p<0.001, two-sample t-test for participants A, B, C), during which their mean RT decreased by 15-40% even though they experienced only 9-12 canceled trials. Reaction time increased gradually as participants experienced a sequence of trials without cancellations (Fig. 6a), which explains why trial cancellations were broadly distributed across sessions and not just limited to the first few sessions of Phase II (Fig. 6b). These results suggest that the participants adjusted their bound height to the environmental statistics throughout the experiment.

**Figure 6.**
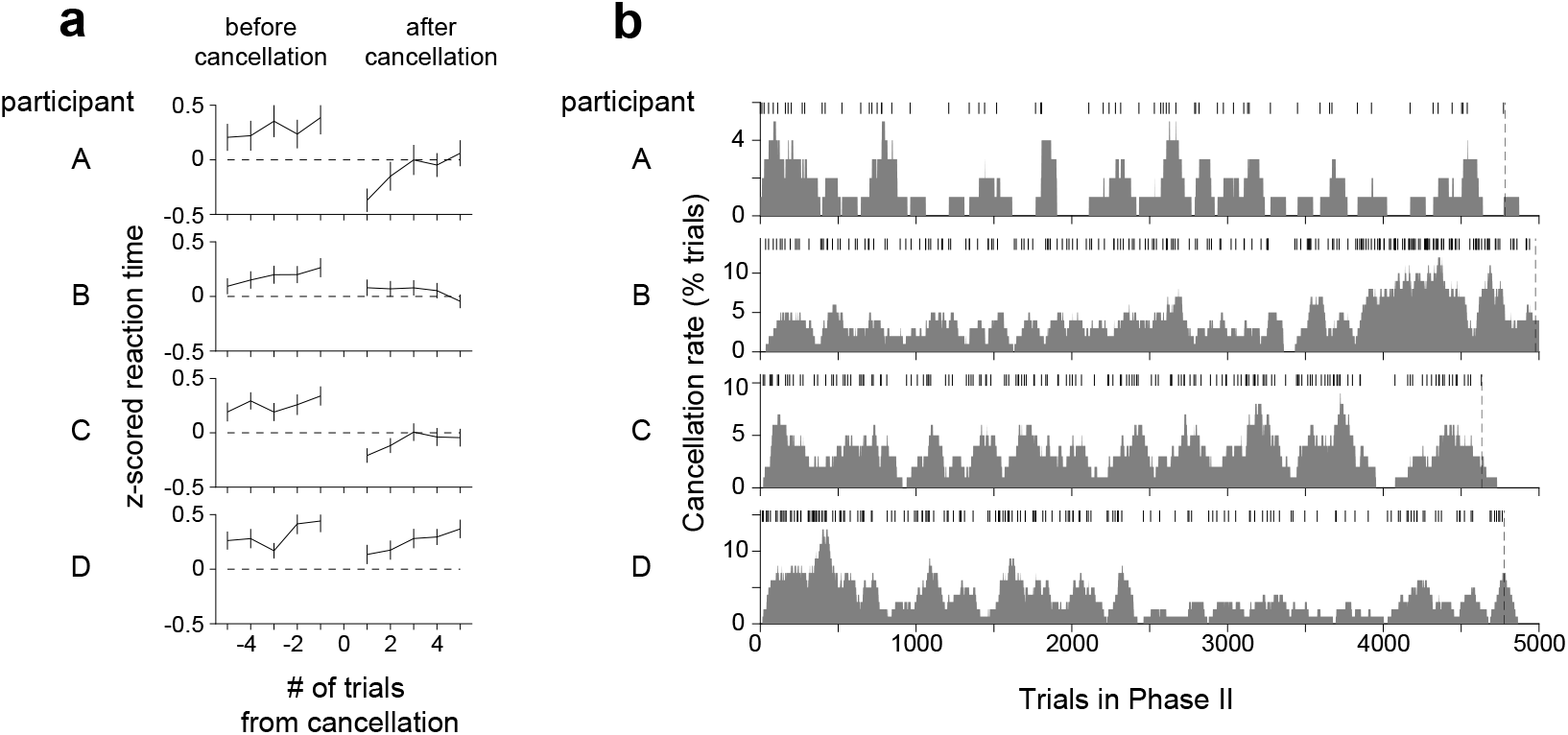
Effect of canceled trials on reaction time. (a) Mean reaction time before and after a canceled trial. Rows correspond to the four participants. RTs from Phase II are z-scored independently for each motion strength, and the mean z-scored RTs are shown for 5 trials before and after the trial cancellation. Error bars indicate s.e.m. Note that the mean z-scored RTs tend to be positive before the cancellation. This is a sign that cancellations are more likely to occur when participants are responding slower than average. (b) Distribution of trial cancellations in Phase II. Rows correspond to the four participants. The ticks indicate trials in which the deadline was reached, resulting in cancellation. The trial cancellation rate (gray area) was calculated using a sliding window of 100 trials and plotted at the last trial in the window (rightmost edge of the window). Vertical dashed lines mark the last trial of Phase II. Panels (a, b) include all trials in Phase II (with and without a provisional deadline).

In the final Phase III of the task, we removed the provisional deadlines to reduce the time cost. Three participants slowed their decisions speed (p<0.01 for participants B, C, D; Fig. 7a), achieving a speed and accuracy that was somewhere between the two phases. They did so by increasing the height of the decision termination bounds relative to those adopted in Phase II (Fig. 7b). However, the height of the bound did not quite reach that of Phase I, even though the experimental design was identical across these two phases. We can only speculate that they still carried a belief, albeit non-conscious, that time can be costly, or perhaps they incorporated the possibility of change in the environment (e.g., volatility). These interpretations aside, the result clearly attests to the flexibility of the time-dependent stopping policy in decision making.

**Figure 7.**
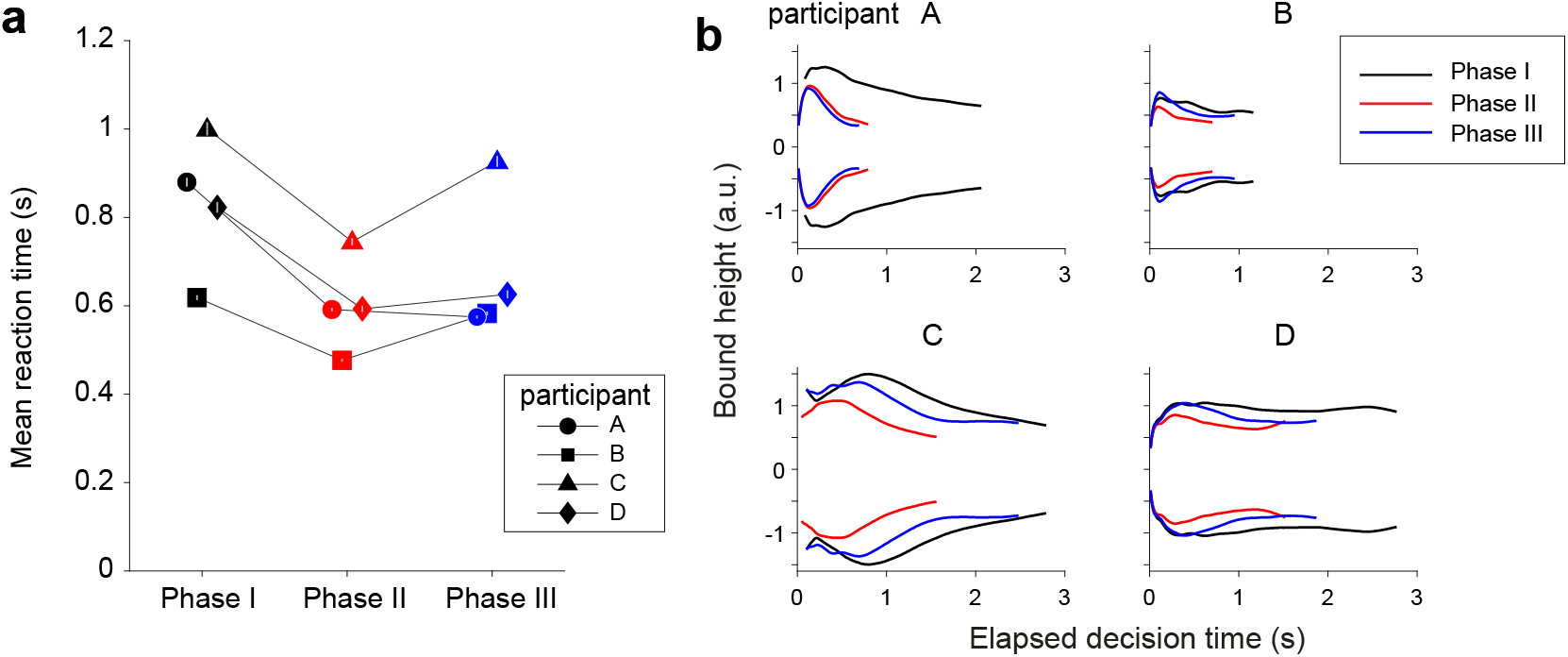
Partial recovery of Phase I bounds after elimination of provisional deadlines. (a) Mean RT across Phases, for each participant (symbols). Trials with a provisional deadline were excluded from the data in Phase II. Symbols and error bars indicate mean ± s.e.m. Color in both panels indicates experimental phase. (b) Shape of the decision-termination bounds derived with the *npb-DDM* for Phases I-III. The bounds are shown for the range corresponding to the 1^st^–99^th^ percentile of decision times for each participant.

## Discussion

We have shown that human observers exercise flexible and refined control over the policy to terminate decisions based on accumulated evidence. For decisions based on a sequence of independent samples of noisy evidence, it is sensible to deliberate until a sufficient quantity of information has accrued. The process may be formalized as a type of sequential sampling of evidence with optional stopping, and with suitable simplifications, it is analogous to bounded drift-diffusion or a biased random walk to bound (Ratcliff, 1978; Smith, 1990). The bounded DDM in particular offers a parsimonious explanation of both decision accuracy and speed.

Perceptual decisions about the direction of dynamic random-dot motion invite the simplifications of the DDM because, on any single trial, samples of evidence are statistically stationary and effectively independent. However, the participant does not know the direction or the difficulty of the stimulus. In many instances, it might make sense to estimate difficulty before deliberating about the question at hand, but owing to the high concentration of difficult conditions in our task, assessing difficulty (i.e., motion strength) would not be fruitful. Specifically, it would not be fruitful to aim for some level of accuracy based on an assessment of motion strength. Indeed participants appear to apply a common criterion to all motion strengths, which leads to the graded accuracy functions in Figure 2b.

The termination rule is an expression of policy in both a mathematical and pragmatic sense. The mathematical sense of policy is borrowed from the field of dynamic programming. By iterating over possible stopping rules, one settles on symmetric bounds, ±B(t), that maximize desiderata. In our experiment, per instruction, this should have been the points earning rate. By pragmatic, we mean that the termination rule is the only important control that the participant can deploy to perform this task. The reliability of the evidence (i.e., signal-to-noise ratio) is controlled by the experimenter and the response of direction-selective neurons in the visual cortex (Shadlen and Newsome, 1994, 1998; Born and Bradley, 2005). A central question in neuroscience concerns the mechanisms responsible for establishing and implementing this bound. Our study contributes to our understanding of the former. It shows that terminating bounds, which operate on the level of accumulated evidence, are dynamic and adjustable, and it shows that they can be adjusted to approximate optimal performance. The latter mechanism of termination has been shown to involve the superior colliculus (Stine et al., 2023).

The tradeoff between speed and accuracy is mediated by control of the threshold, or bounds, for terminating a decision. In the standard DDM, these bounds are assumed to be stationary, that is, time-independent or flat. However, this simplification, inspired by Wald’s sequential probability ratio test (Wald, 1947; Wald and Wolfowitz, 1948), would predict that the time-dependent accuracy functions ought to be flat. Clearly they are not flat, but decline as a function of reaction time (Fig. 2c-d). Such collapsing bounds also explain the longer RTs accompanying errors (Fig. 2a) and the shapes of RT distributions (Fig. S1). Indeed, collapsing bounds are the natural extension of Wald’s normative theory for the common situation in which a variety of possible difficulties might be encountered (Drugowitsch et al., 2012). The key intuition is that as time elapses, a decision that has yet to terminate is more likely to be based on evidence from a less reliable source (i.e., lower signal to noise) and hence more likely to result in an error. Consequently, the deliberation time is progressively more costly. Collapsing bounds might also incorporate other costs associated with deliberation time, such as mental effort (e.g., allocation of attentional resources).

The manipulation introduced in Phase II of our experiment provides a direct test of these ideas. It shows that the bounds change shape in response to a subtle change in experimental conditions. We argue that the shape of the bounds in Phases I and III of the experiment also reflect the cost of decision time, but we did not test this directly, for example, by changing the point system. Our manipulation led to shorter RTs, and they did so in a way that is most consistent with linear scaling of the bounds. The essential feature of this mechanism is that decisions terminate based on the state of the evidence.

An alternative to a change in bounds is that the intervention applied in Phase II made participants more attentive. This could speed up decisions if a change in attention were to reduce noise of the sensory representation, as is commonly believed (McAdams and Maunsell, 1999; Bichot et al., 2005; Cohen and Maunsell, 2009). However, this mechanism would not explain the change in RT at the lowest coherences. In fact, lowering the noise at the lowest coherences would increase RT because those decision times are dominated by diffusion, not drift (Zylberberg et al., 2016). Supporting this interpretation, we found that the signal-to-noise parameter (*κ*) explained only a small fraction of the reduction in RT from Phase I to Phase II (Fig. S3).

The mechanism at play in all three phases of the experiment helps the participant maximize their points earning rate. We estimated this optimal rate using dynamic programming. After an adjustment period, participants came within 99% of the optimal rate. We are loath to read too much into this small discrepancy. The process is, after all, confounded by perceptual learning and other adjustments (e.g., to the eye tracker). Moreover, the estimate of optimal performance assumes perfect knowledge of the prior over stimulus-difficulty and the temporal structure of the task, whereas participants can only approximate this. Finally, although the definition of optimal is clear, it is hard to know what participants actually experience. For example, it has been argued that mental effort is itself costly (Drugowitsch et al., 2012).

Our non-parametric method for finding decision bounds allowed us to identify features of their shape that could have been overlooked by more conventional methods based on parametric forms. Using the *npb-DDM*, we found that the shape of the bounds was highly consistent across different phases of the task. This consistency was revealed by an analysis in which we used simple transformations—linear scaling in time and magnitude—applied to the bounds obtained from Phase I to explain the data of Phase II. The behavioral data in Phase II was best explained by scaling the bounds obtained from the same (rather than different) participant. This result suggests that bound shape is idiosyncratic. This idiosyncracy is not expected from the solution of the optimal model, which produced similar bounds across participants. However, the notion that the bound shape is an individual ‘fingerprint’ should be considered preliminary, since the experiment was performed by a small number of participants.

People (and other animals) often take longer to make a choice when the decision is preceded by another one in which they chose incorrectly, a phenomenon known as post-error slowing (PES) (Rabbitt, 1966). This increase in reaction time is due to an increase in the amount of evidence required to make a choice, mediated by a change in the height of the decision-termination bounds (Purcell and Kiani, 2016). In our experiment, the cancellation of a trial also leads to an adjustment in response speed, but in the opposite direction to that observed in the PES. That is, people were faster immediately after experiencing a cancellation (Fig. 6a). This distinction suggests that the adjustment of decision policy after an error requires a causal inference about the origin of the error: insufficient urgency versus a lax stopping criterion. We imagine that the degree of confidence in the evolving but canceled decision might inform this assignment, but this remains to be seen. At the very least, the distinction attests to the flexibility of the decision-making process, and the possibility of precisely and rapidly altering the parameters that control the speed-accuracy regime.

Our study does not address the neural mechanisms underlying the establishment and implementation of the bound, but studies in non-human primates provide insight. When monkeys perform the direction discrimination task, they also exhibit the same regularities as our human participants, including declining time-dependent accuracy, slow errors, and RT distributions suggestive of collapsing decision bounds (Drugowitsch et al., 2012; Kira et al., 2015). However, the neural implementation is not a change in a threshold per se. Neurons in several brain areas are known to represent the accumulation of noisy evidence resembling an integral of the difference in signals from left- and right-preferring direction-selective neurons in areas MT/MST. The activity of neurons in LIP gives a clear indication of a terminating threshold—a point of minimum variance just before the response (Roitman and Shadlen, 2002; Churchland et al., 2011; Kira et al., 2015)—implying that a downstream area terminates the decision when the firing rate reaches a critical level. However, this threshold is not time-dependent. Instead, the DDM is realized by a race between accumulators for left (and against right) and for right (and against left) (Mazurek et al., 2003; Gold and Shadlen, 2007; Churchland et al., 2008; Kira et al., 2015), such that the first accumulation to reach a positive termination bound establishes the choice. In this architecture, a change in bound height is equivalent to a change in starting point for both processes, and a collapsing bound can be implemented as a time-dependent signal that is added to both accumulators, what we term an urgency signal. This urgency signal is evident in the neural recordings from area LIP (Churchland et al., 2008; Kira et al., 2015) and it changes if the monkey changes the speed-accuracy regime (Hanks et al., 2014).

The implication is that setting B(t) and detecting a bound crossing are likely to involve different structures. The first involves the generation of a time-dependent urgency signal that is added to the neural representation of cumulative noise and signal (Janssen and Shadlen, 2005; Churchland et al., 2008; Hanks et al., 2014; Kira et al., 2015), whereas the latter is a simpler threshold crossing detection that operates on the firing rates of these neurons. It can be carried out by a downstream structure, such as the superior colliculus, in a manner that is consistent regardless of time costs (Stine et al., 2023). There is something to be said for the parsimony of this design. Whereas the establishment of B(t) appears to accommodate detailed policy considerations, the termination threshold appears to operate generically, that is to say, independent of these considerations.

## Methods

### Participants

Participants were four young adults (3 males and 1 female) with normal or corrected-to-normal visual acuity. All the participants participated in the experiment with informed written consent. They either volunteered or were paid $15 per hour during the experiment. All experimental procedures were approved by the institutional review board at the University of Washington.

### Apparatus

Participants were seated in front of the display. A chin rest and forehead bar were used to stabilize the head of the participant for the duration of each experimental session. The stimuli were displayed on a flat-screen CRT video monitor (19-in View Sonic PF790, 800 by 600 pixels, viewing distance = 57 cm, subtending 35.0°by 26.6°with 22.1 pixel/deg at screen center; refresh rate = 75 Hz) controlled by a Macintosh G5 (2×2.8GHz, Mac OS 10.5 with an ATI Radeon HD 2600 graphics card). Stimuli were generated using the Psychophysics Toolbox Version 2 (Brainard, 1997; Pelli, 1997) for MATLAB (Version 7.5, Mathworks). Eye movements in one eye were recorded with a 1000 Hz sampling rate by a noninvasive infrared video-tracking system (EyeLink 1000, SR Research).

### Motion direction discrimination task

Participants discriminated the direction of dynamic random-dot motion in a choice-reaction time paradigm (Fig. 1a). Trials began with the appearance of a fixation spot (0.3°diameter) in the center of the screen. Participants were required to look at the fixation spot and maintain their fixation within 4°x4°of their initial fixation. Once fixation was attained, two red targets (0.5°diameter) were presented 6°to the left and right of the fixation spot. After a variable delay (200–500ms, exponentially distributed; mean 300ms), the random dot motion stimulus appeared within a 5°circular aperture centered on the fixation spot. Dots were 2 by 2 pixels (0.09°square), with a density of 16.7 dots/deg^2^/s. On each trial, the motion coherence was selected randomly from the list (±0, ±3.2, ±6.4, ±12.8, ±25.6, ±51.2%). The absolute value of the motion coherence—referred to as the *motion strength*—specifies the fraction of dots that are displaced in motion as opposed to randomly repositioned. The sign of the motion coherence indicates the direction of motion (negative for leftward and positive for rightward).

The displacement of the coherently moving dots was such that the apparent speed of motion was 5 dva⋅s−1 (dva: degrees visual angle). Details on the generation of the random dot motion have been described in previous studies (Roitman and Shadlen, 2002; Palmer et al., 2005).

After the motion stimulus onset, participants were required to decide the direction of the motion stimulus and make a saccadic eye movement to the corresponding choice targets. When a saccade was detected within 6°x6°around the chosen target, the trial was classified as either correct or error based on whether a chosen target direction and an assigned motion direction were the same or opposite, respectively. In these trials, reaction time (RT) was defined as the time from the motion stimulus onset to the initiation of the saccade, which was detected when the gaze first exited the 4°x4°fixation window. When participants broke a fixation prematurely (e.g., eye blinks) or made an invalid saccade, these trials were terminated immediately and labeled as “*aborted* “.

Each trial was followed by a 2 s inter-trial interval. Participants performed trials repeatedly during a 13 min session (234.3 ± 11.0 trials/session, mean ± s.d., n = 160 sessions from 4 participants) and completed two sessions a day with a short break (5–10min) in between.

### Scoring system

Participants earned one point for a correct choice and lost one point for an erroneous choice. The designation of correct or error was random for the 0% coherence trials. The total point did not change for aborted trials of any type (blinks, fixation inaccuracy, etc.). During the 2 s of an inter-trial interval, participants received visual and auditory feedback about their performance. The visual feedback indicated the total points in digits and the earned points per minute (points earning rate) in a bar graph, which included the current points earning rate during the latest 2 min and a history of the points earning rate for every 2 min from the beginning of a session (Fig. 1a). The participants were instructed to maximize their points earning rate during a session.

### Experimental schedule

We divided the experiment into three phases (Fig. 1b). We did not explain the experimental schedule to the participants, so that the prior knowledge of the task would not bias their behavior.

In Phase I, we measured the baseline performance. The participants were allowed to make a decision whenever they were ready within 5 s from the motion stimulus onset. These trials are referred to as *standard trials*. Because participants never viewed the stimulus for the whole duration, participants performed the task as if there were no time limit. While the participants performed the task, we monitored their performance by tracking the total number of points earned and the earning rate (points per min). We also tracked the mean RT on the 0% coherence trials (participants A and B only). The participants performed the task repeatedly until their performance was deemed stable in terms of RT and points earning rate for at least 10 sessions. We use the last 10 sessions (2154–2389 trials) to characterize the baseline level of performance.

In Phase II (20 sessions), we evaluated the effect of increased time cost by introducing provisional deadlines. The deadline trials and standard trials were randomly interleaved during this phase. If the participant made a decision before a deadline, the trial ended in the same way as in the standard trial. If participants had not made a decision by this deadline, however, the trial was terminated immediately as if they broke a fixation, and labeled as “canceled”. Each provisional deadline was randomly drawn from a time-shifted Rayleigh distribution:

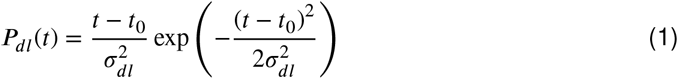

where *t*_*0*_ is a time shift that sets the minimum time for the deadline, and *σ*_*dl*_ is the scale parameter of the distribution. These parameters were tailored for each participant based on his or her RT distributions measured in Phase I (Table 1). Specifically, the parameters were chosen to cancel ∼10% of the trials in Phase II, had the participants performed with the same decision speed as in Phase I. Note that the mean RT for deadline trials is expected to be shorter than RT for standard trials because RT has to be shorter than a deadline in each trial. To eliminate the confound that this attrition accounts for the shorter RT in Phase II than in Phase I, deadline trials were excluded and only standard trials were used for the analyses, except for those in Figure 6.

In Phase III (10 sessions), we evaluated a washout effect by eliminating the provisional deadlines. The participants performed standard trials as in Phase I.

### Statistical analysis of choice and RT data

We used linear regression to test the statistical significance of the difference in RT between phases:

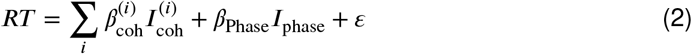

where the motion strength of each trial (0, 3.2, 6.4, 12.8, 25.6 or 51.2%) is indexed by the indicator variable 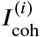. *I*_Phase_ is an indicator variable for phases, which takes 1 for one phase and 0 for another phase in comparison. *ε* represents a Gaussian error. We used *glmfit*.*m* in MATLAB (MathWorks) for fitting the above model separately to correct and incorrect choices, and testing whether *β*_Phase_ (RT difference between phases) is significantly different from zero by t-test using the s.e. of the estimated regression coefficient.

We used logistic regression to determine if there is a significant difference in accuracy between Phase I and Phase II. The model that was fit separately for each participant is:

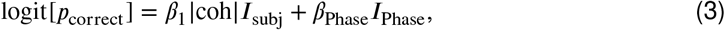

and the model fit combining data across participants is:

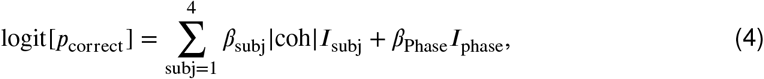

where *I*_subj_ is an indicator variable that takes a value of 1 for trials completed by participant subj and zero otherwise, and *I*_phase_ is an indicator variable that takes a value of 1 for Phase II trials and 0 otherwise. We used a likelihood-ratio test for nested models with and without the *β*_Phase_ term to evaluate the null hypothesis that *β*_Phase_ is equal to zero.

### Bounded drift-diffusion model

We analyzed participants’ performance in the framework of a bounded drift-diffusion model (Fig. 3a). In this model, the decision variable (DV), denoted by *x*, evolves over time by following a Wiener process with a constant drift:

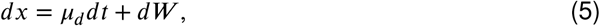

where *W* is a Wiener process or Brownian motion. The drift term *μ*_*d*_ is proportional to the motion strength: *μ*_*d*_ = *κ* ⋅ coh, where coh is the motion coherence with a positive or negative sign for rightward or leftward motion, respectively. *κ* is a free parameter that determines a signal-to-noise ratio in our model. Together, when the motion coherence is rightward (or leftward), it is likely to furnish positive (or negative) momentary evidence. The stochastic process produces the decision variable, *x*(*t*), continuously evolving while −*B*(*t*) < *x*(*t*) < *B*(*t*), where ±*B*(*t*) are the upper and lower terminating bounds for right and left choices, respectively. Note that these bounds are symmetric about *x* = 0, but they are not assumed to be time-invariant (i.e., ‘flat’). The first passage time determines the choice and the decision time (*T*_*d*_). The reaction time (saccade initiation) relative to motion onset includes additional sensory and motor delays, which we assume to be independent of motion strength and direction and refer to as the non-decision time (*T*_*nd*_). The *T*_*nd*_ on each trial is realized as a normally distributed random value, 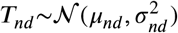. We fit *μ*_*nd*_ and 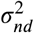 independently for each participant and experimental phase.

### Nonparametric-bound Drift-Diffusion Model (*npb-DDM*)

We developed a non-parametric method for estimating the shape of the decision terminating bounds that does not require assuming a functional form.

The key intuition is that in the bounded drift-diffusion model, if the drift rate is known, then the distribution of decision times is uniquely determined by the time course of the bounds, ±*B*(*t*). Given this unique relation, the process can be inverted to infer *B*(*t*) from the empirical distribution of decision times.

Because of the non-decision latencies, the distribution of decision times is not directly observed; yet, we can approximate it from the observed reaction times. The decision time for each trial *i* is estimated by 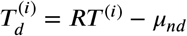, where *μ*_*nd*_ is the mean non-decision time. The probability density function for the decision times, 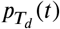, is calculated from the single-trial decision times, 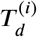, using non-parametric kernel smoothing with an Epanechnikov kernel (inverted parabola with s.d. = 0.1 s).

After estimating 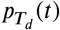, we alternated two operations, *propagation* and *match*, iteratively over time.

#### Propagation

In this step, we propagate the probability distribution of the decision variable for a single time step. The probability mass was propagated by numerically solving the Fokker-Planck equation that describes the dynamics of a Wiener process with a constant drift:

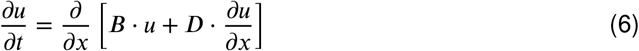

where *B* = −*μ*_*d*_ (≡ −*κ* ⋅ coh), and *D* = *σ*^2^⁄2. The single-particle distribution function in the space ⟨*x, t*⟩ is *u* (*x, t*). Each motion coherence is treated independently. For numerical stability and to avoid negative probability mass that results from semi-implicit methods when the initial condition is defined by a *δ*-function, we used a fully implicit numerical scheme (Chang and Cooper, 1970).

After solving the Fokker-Plank equation for a single time step, we obtained *p*(*x, t*| coh), which specifies the probability that the DV takes the value *x* at time *t* given that the underlying coherence is coh. We then marginalize over motion coherences to obtain the probability distribution of the DV at time *t* as:

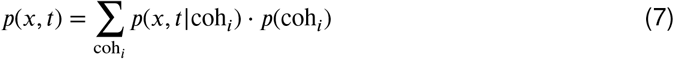

#### Match

Since we approximated the distribution of decision times, 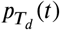, we know the probability of bound-crossing at time *t*. Since there is a monotonic relation between the bound height and the proportion of DV that crossed the bound, we can set the height of the symmetric bounds ±*B*(*t*) such that the probability mass that is absorbed at either bound at time *t*,

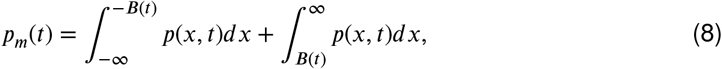

is equal to the probability that the decision terminated at time *t* as obtained from the empirical distribution of decision times, *pT d* (*t*). *B*(*t*) was determined at each time step by root finding, bracketing B within a sequence of diminishing intervals. The steps of *propagation* and *match* were repeated iteratively over time in steps of *dt* = 0.5 ms.

#### Fitting the model’s free parameters

Given the set of model’s free parameters *θ* = [*κ, μ*_*nd*_, *σ*_*nd*_] and the non-parametric decision bounds derived as above, the model can predict the joint probability distribution of the RT and choice direction under each motion coherence, *P* (RT, choice | *θ*, coh). For the experimental data, *D*, comprising *RT* s and choice directions across individual trials in each Phase, we searched for a set of parameters that maximizes the likelihood of observing the data:

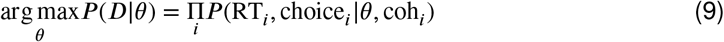

where *i* indexes individual trials for a specific participant and phase.

### Cross-fitting of scaled bounds

Here we describe the statistical analysis bearing on our claim that participants apply a scaled version of their unique bound shape across the phases of the experiment. For each participant and Phase, we fit the *npb-DDM* to the data, 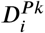, where the subscript *i* ∈ {1..4} indexes the participants and the superscript *P k* (*P* 1 or *P* 2) indexes Phases. This yields fit-parameters 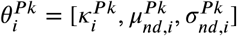, and bounds 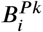. We ask how well the Phase II data from participant *i* can be explained by linear scaling of the bounds that were obtained from Phase I data from participant *j*. The parameters of the DDM (signal-to-noise and non-decision time) were set to those obtained by fitting the *npb-DDM* to Phase II data from participant 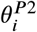. We use two free parameters, **s** = {*st, sm*}, to scale the bounds as a function of time and magnitude:

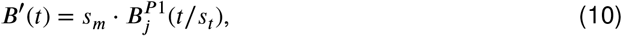

and fit **s** using maximum log-likelihood:

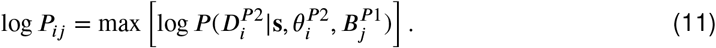

The quality of the fit was quantified as the difference between the log likelihood obtained from the best-fitting scaled-bound model and the *npb-DDM*:

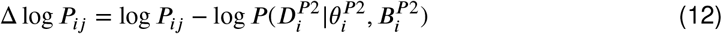

If Phase II data of participant *i* are best explained by scaling of Phase I bounds from the same participant 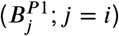 than other participants (*j* ≠ *i*), then Δ log *P*_*ij*_ should be greatest when *j* = *i*. To adjust for the difference in the number of trials across participants in Phase II, Figure 5b shows the average log likelihood across data points (trials): 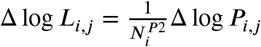, where 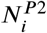 is the number of trials for participant *i* in Phase II.

We use a permutation analysis to test the hypothesis that choices and RTs from Phase II are best explained by scaling the decision bounds for Phase I from the same participant, against the alternative that scaling from the same or different participants is inconsequential. For each participant *i*, we rank-ordered Δ log *P*_*i*\*_, and created a 4-by-4 matrix of the rank order. If each participant’s data were best explained by scaling his or her own bounds, then the diagonal elements of the matrix should be [1, 1, 1, 1], which sums to 4. In the data, the ranks were [1, 2, 1, 1], which sums to 5. There are 4 patterns (permutations) of these ranks that sum to 5. In contrast, under the null hypothesis, { ℋ_0_: the rank order is random}, there are 4^4^ (= 256) possible patterns of the ranks across the diagonal elements. Therefore, the p-value that the sum of the ranks is 5 or less, as in our results, is (1+4)/256 (*p* = 0.020).

### Derivation of the optimal decision strategy

A decision policy maps a state of accumulated evidence and elapsed time to an action. The possible actions are to accumulate more information or to stop the decision process and make a left or right choice. Deriving the optimal decision policy is not trivial since it depends on the opportunity cost of time, which itself depends on the decision policy. We formulate the decision process as a Markov decision process (MDP) and use dynamic programming to derive the optimal policy (i.e., the one that maximizes points earned per unit of time) in each participant and phase of our experiment.

A Goal MDP is described by: a non-empty state space *S*, an initial state *s*_0_, a goal state *s*_*g*_, a set of actions *A*(*s*) executable in state *s*, positive or negative reward *r*(*a, s*) obtained after performing action *a* in state *s*, and transition probabilities *P*_*a*_ (*s*^′^ *s*) representing the probability of transitioning to state *s*^′^ after performing action *a* in state *s* (Geffner and Bonet, 2013).

The states *s* ∈ *S* are defined by a tuple *s* = ⟨*x, t* ⟩, where *x* is the accumulated motion evidence in favor of the rightward target, and *t* is the elapsed decision time. At time zero, there is no accumulated evidence favoring either choice, *s*_0_ = ⟨*x* = 0, *t* = 0⟩. Three actions are applicable in each state: terminating the trial by making a left choice (left) or a right choice (right), or accumulating more evidence by maintaining the eye fixation (fix).

Choosing a terminal action (left or right) leads to a cost-free absorbing goal state. In contrast, state transitions that follow the fixation (fix) are stochastic because of the noise in the momentary evidence even if the motion coherence was known. We discretize time and the states of accumulated evidence in small steps of *δt* and *δx*, respectively.

The transition probability *p*_*fix*_ (*s*^′^|*s*) specifies the likelihood of transitioning from state *s* = ⟨*x, t*⟩to state *s* = ⟨*x*’, *t*’⟩ after selecting action fix in state *s*. Because time is discretized in steps of *δt*, the only transitions with non-zero probability are those to states *s*^′^ in which *t*^′^ = *t* + *δt*.

The transition probabilities depend on motion coherence, which is unknown to the decision maker. Thus, the optimal decision maker must marginalize over the motion coherences to calculate *P*_*f ix*_ (*s*^′^|*s*), as follows:

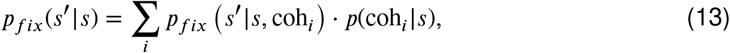

where *p*_*f ix* (_*s*^’^|*s*, coh_*i*_)is the transition probability from state *s* to state *s*^′^ if the motion coherence were coh_*i*_. As in the drift-diffusion model, we assume that the evidence gathered in a small time step *δt* follows a normal distribution with a mean equal to *κ* ⋅ coh_*i*_ ⋅ *δt*, and the variance equal to *δt*, thus:

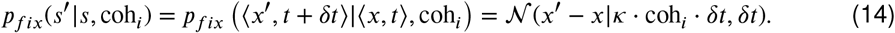

The transition probability *P*_*fix*_ (*s*^′^|*s*) also depends on *p*(coh_*i*_|*s*), the probability that motion coherence is coh_*i*_ when in state *s* = ⟨*x,t*⟩ (Eq. 13). This probability can be calculated by Bayes’ rule. Assuming that all motion coherences have equal prior probability (as in the experiment):

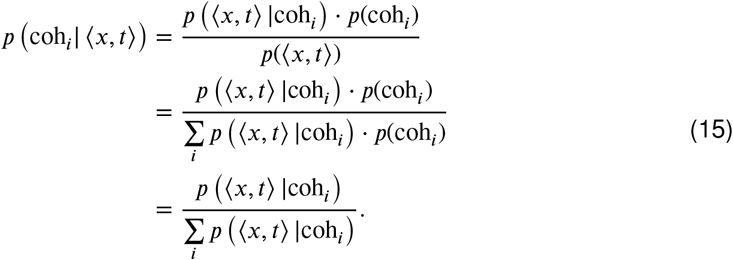

If there are no decision termination bounds,

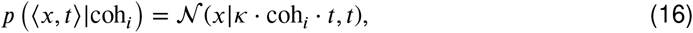

and thus:

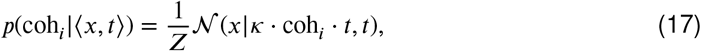

where *Z* is a normalization constant which assures that the sum of *p* (coh_*i*_ ⟨*x, t* ⟩) over all coherences adds to one.

Following the approach of Drugowitsch et al. (2012), we show that Eq. 17 still holds in the presence of termination bounds and provisional deadlines. Let 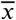 be a trajectory that leads Ω(*x, t*) denote the set of all 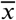 leading to ⟨ *x, t*⟩ without reaching the termination bounds. With termination bounds and provisional deadlines,

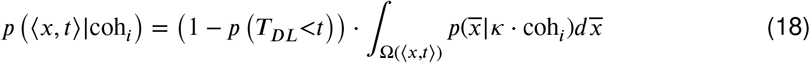

where *p T*_*DL*_<*t* is the probability that a deadline occurs before time *t*, and 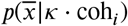 is the probability of a single trajectory 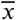, which can be expanded across time steps as follows,

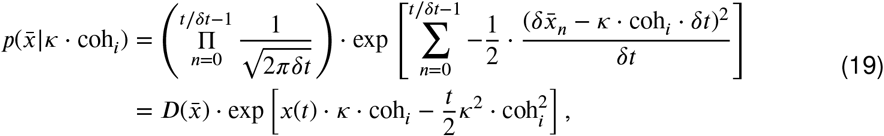

where 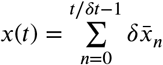 and 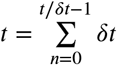 were used to substitute the sums. 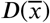 is a function of the trajectory 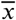, and is independent of the motion coherence. That is, for each trajectory, there is a coherence-independent scaler 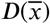, such that Equation 18 becomes,

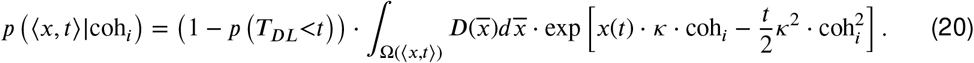

In Eq. 20, the coherence-dependent exponential term is multiplied by the coherence-independent terms that depends on the bound shape and the provisional deadline. Consequently, for any two motion coherences, coh_*i*_ and coh_*j*_, the ratio *p* (⟨*x, t*⟩| coh_*i*_) /*p* (⟨*x, t*⟩|coh_*j*_) is independent of the bound shape and the presence of a provisional deadline, and equal to the ratio of two normal distributions indexed by Eq. 16. Therefore, Eq. 17 holds independently of the bound shape and the presence or absence of provisional deadlines.

To derive the policy that maximizes the points earning rate, we must take into account the opportunity cost associated with the passage of time. That is, while prolonging deliberation increases the decision accuracy on average, deliberating for too long will decrease the overall earning rate. Using established methods (Bertsekas, 1995), we find the decision policy that maximizes the earning rate. For phases I and III (no deadlines), the Bellman equation is:

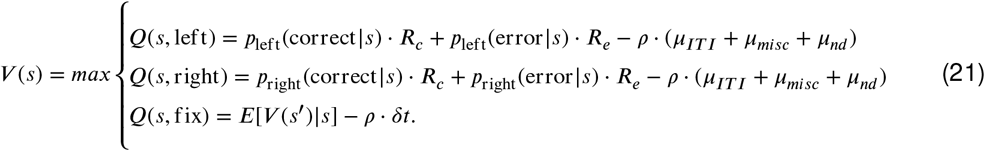

*Q*(*s, a*) is the state-action value function representing the reward expected for performing action *a* in state *s. p*_*a*_ (correct |*s*) is the probability of being correct after taking action *a* in state *s*, and *p*_*a*_ (error| *s*) is calculated as 1 − *p*_a_ (correct *s*). Parameter *μ*_*nd*_ is the mean non-decision time, and *μ*_*IT I*_ is the mean inter-trial interval. *μ*_*misc*_ is a mean time for miscellaneous experimental latencies, including the time to acquire fixation, the delay after the fixation to the motion onset, and the validation time for a saccadic response. *R*_*c*_ and *R*_*e*_ are the rewards (points) obtained after a correct and error choice, respectively. The expectation in *Q*(*s*, fix) is an expectation over all future states *s*^′^ that result from being in *s* and accumulate evidence for an additional *δt*. The earning rate, *ρ*, is unknown because it depends on the decision policy itself. We searched for the value of *ρ* iteratively by root finding, using the bisection method (Bertsekas, 1995; Drugowitsch et al., 2012).

To derive the optimal policy for Phase II, we modified the equation for fixation by including the cost of trial cancellations due to deadlines:

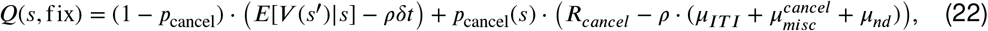

where *R*_*cancel*_ is the reward (points) following a trial cancellation. *p*_*cancel*_(*s*) is the probability that a deadline occurs at time *t* given that it has not occurred before. If all trials have a provisional deadline, then *p*_*cancel*_(*s*) is given by the hazard function of the Rayleigh distribution. However, because trials with and without provisional deadlines are randomly interleaved, *p*_*cancel*_(*s*) also depends on the probability that the trial has a provisional deadline (*d*^+^) given that time *t* has elapsed without a cancellation, *p*(*d*^+^|*t*):

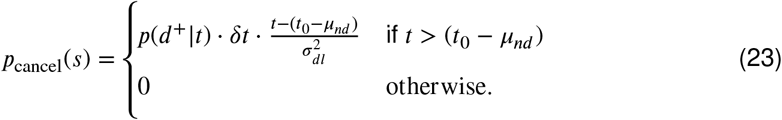

The probability that the trial has a deadline given that no deadline has been reached by time *t*, |*p*(*d*^+^ |*t*), can be calculated by Bayes’ rule:

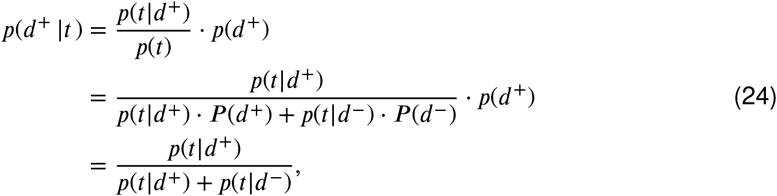

where *p*(*t*| *d*^+^) and *p*(*t*| *d*^−^) are the probabilities that time *t* has elapsed without a cancellation on trials with and without a provisional deadline, respectively. *p*(*t* |*d*^−^) is equal to 1 because there are no cancellations due to provisional deadlines in *d*^−^ trials. *p*(*t*| *d*^+^) is given by the survival function for the Rayleigh distribution:

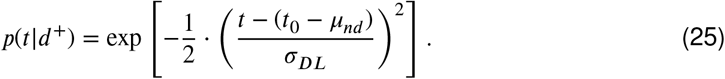

The optimal policies depicted in Figures 4 and 7 were derived with the following parameter values: *R*_*c*_ = +1 [point], *R*_*e*_ = −1 [point], *R*_*cancel*_ = 0 [point], *δt* = 0.5 [ms], *δx* = 0.02 [a.u.], *μ*_*IT I*_ = 2.0 [s], *μ*_*misc*_ = 0.7 [s], 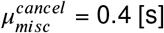 (note *μ*_*misc*_ is longer for non-canceled trials due to the saccade validation time), and fitted parameters (*κ* and *μnd*) for each participant and phase (Table 2).

**Table 2.** Fitted model parameters for the *npb-DDM*.

| participant | Phase | Fitted parameters of the empirical bound model |  |  |
| --- | --- | --- | --- | --- |
| | | $K'$:<br>scaling factor for SNR | $\mu_{nd}$:<br>mean of non-decision time (s) | $\sigma_{nd}$:<br>s.d. of non-decision time (s) |
| A | I | 16.19 | 0.27 | 0.001 |
|  | II | 20.53 | 0.30 | 0.036 |
|  | III | 20.91 | 0.32 | 0.053 |
| B | I | 16.93 | 0.31 | 0.022 |
|  | II | 16.26 | 0.27 | 0.027 |
|  | III | 20.39 | 0.32 | 0.077 |
| C | I | 14.93 | 0.19 | 0.014 |
|  | II | 16.24 | 0.25 | 0.001 |
|  | III | 14.92 | 0.19 | 0.002 |
| D | I | 20.85 | 0.32 | 0.027 |
|  | II | 22.89 | 0.28 | 0.035 |
|  | III | 28.17 | 0.24 | 0.040 |

## Author contributions

SK and MNS designed the study. SK developed the task and performed the experiment. SK and AZ analyzed the data. AZ conceived and designed the *npb-DDM* and the optimal model. SK wrote the first draft of the manuscript. All authors revised the manuscript.

## Acknowledgments

We thank NaYoung So and Natalie Steinemann for their comments on the manuscript. This work was supported by the National Institutes of Health (R01NS113113 to M.N.S.), the Air Force Office of Scientific Research under award (FA9550-22-1-0337 to M.N.S) and the Howard Hughes Medical Institute (M.N.S.).

## Supplementary information

**Figure S1.**
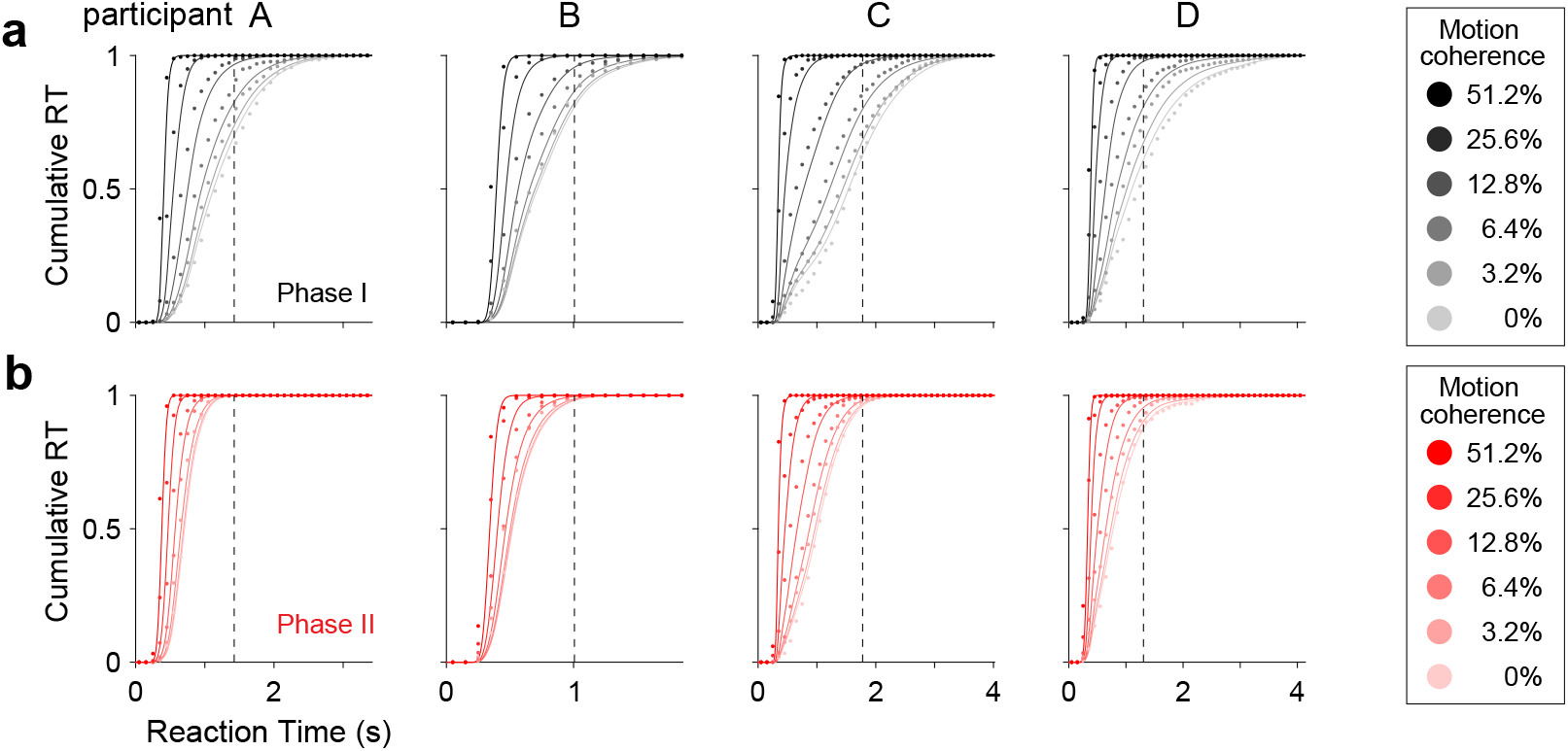
Reaction-time distribution across motion strengths predicted by *npb-DDM*. Cumulative distribution of observed RT (circles, 100 ms bins) and RT predicted by *npb-DDM* (curves) in Phase I (a) and II (b). Trials with a provisional deadline were excluded from the data in Phase II. Vertical dashed lines show mean provisional cancellation time for each participant, calculated from Phase II trials with provisional deadlines.

**Figure S2.**
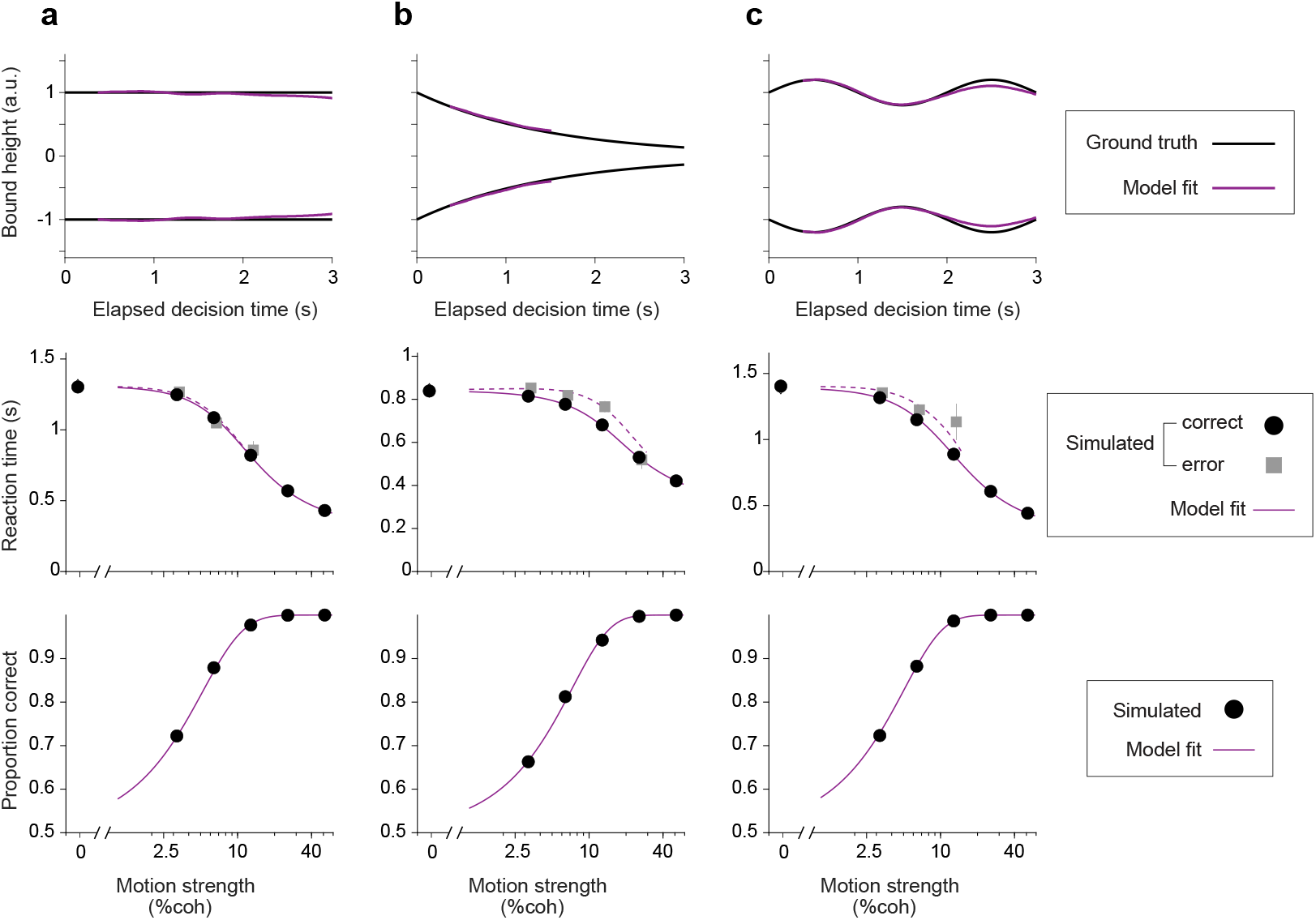
Validation of the *npb-DDM*. (a) We simulated 1,000 trials per motion coherence of a drift-diffusion model with the decision-termination bounds depicted by the black curves in the top panel. Mean reaction time (middle panel) and proportion correct (bottom panel) are shown as a function of motion strength (circles and squares). Red curves were obtained from fits of the *npb-DDM*. (b-c) Same as panel (a) but for decision-termination bounds that decay exponentially (b) or oscillate (c). In all cases, the *npb-DDM* was able to recover the true shape of the bounds and fit the choice and RT data.

**Figure S3.**
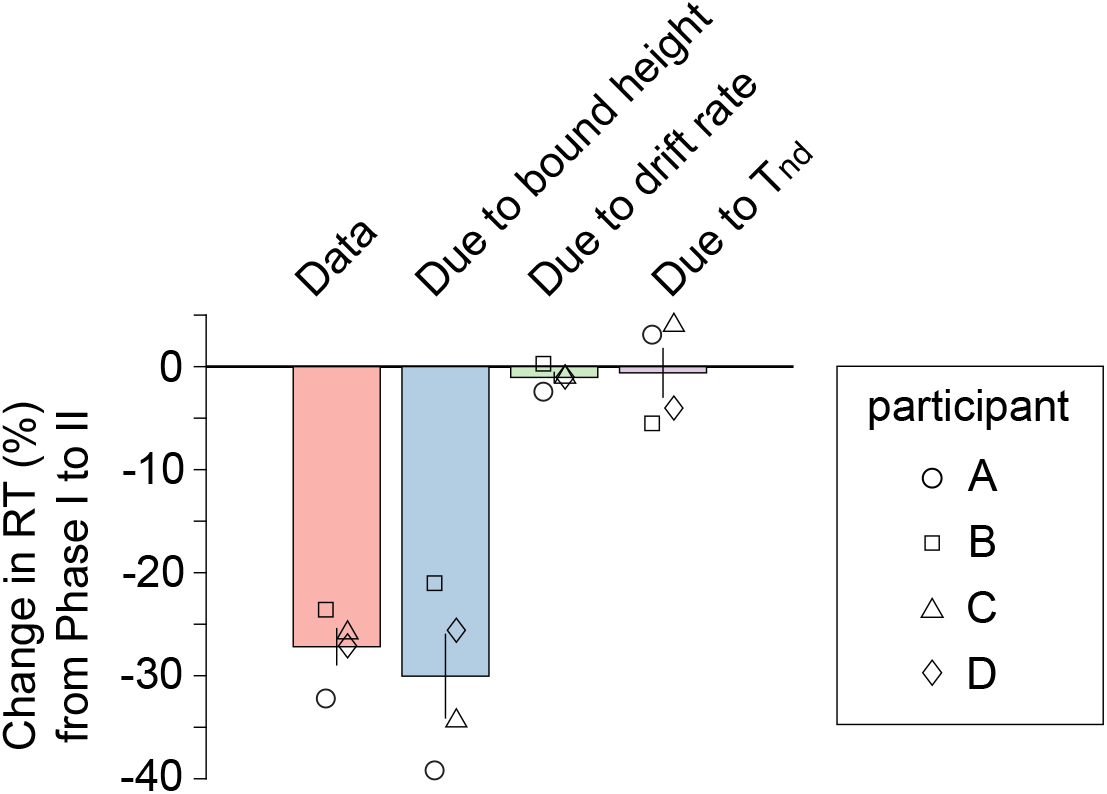
RT reduction due to the change in each model parameter from Phase I to II. Reduction in mean RT (%) from Phase I to II. Bars indicate mean across participants. Individual participant data are indicated by the open symbols. The leftmost (red) bar corresponds to the experimentally observed change in RT from Phase I to II. The next three bars correspond to model expectations. These are obtained from numerical solutions of the *npb-DDM* with parameters fit to the Phase I data, except for one of the parameters that is replaced with the value obtained from fitting Phase II data. The replaced parameters are bound height (blue bar), drift rate (green), and mean non-decision time (purple). The reductions in mean RT are significantly different from 0% for the changes due to bound height (p=0.0027; one-tailed t-test), but not for the changes due to drift rate (p = 0.078) or *T*_*nd*_(p = 0.41).

## Notes

### Competing Interest Statement

The authors have declared no competing interest.

